# Younger adult brain utilizes interhemispheric strategy of recruiting ipsilateral dorsal premotor cortex for complex finger movement, but not aging brain

**DOI:** 10.1101/2024.05.26.595953

**Authors:** Gen Miura, Tomoyo Morita, Jihoon Park, Eiichi Naito

## Abstract

The ipsilateral sensorimotor cortices (dorsal premotor cortex [PMd], primary motor cortex [M1], primary somatosensory cortex [S1], and superior parietal cortex of Area 2) are often activated when the healthy younger and older adult brains perform complex finger movements. Prompted by clinical evidence that the ipsilateral PMd plays particularly important roles to complement the movements after stroke, we tested whether the ipsilateral PMd also plays particularly important (complementary) roles among the ipsilateral sensorimotor cortices when healthy younger adults perform a complex motor task by augmenting interhemispheric functional connectivity with the contralateral sensorimotor cortices. We also examined possible strategic difference when older adults with degraded interhemispheric inhibition perform the complex task when compared to younger adults with mature one.

We addressed these questions by measuring brain activity with functional magnetic resonance imaging while healthy right-handed younger and older adults performed simple (button pressing with the right index finger) and complex (stick rotation requiring coordination between the right fingers) motor tasks.

In younger group, the ipsilateral PMd, S1, Area 2 activated during the complex task, while the ipsilateral M1 remained deactivated as in the simple task, suggesting an existence of interregional difference. The ipsilateral PMd and S1/Area 2 more activated in the individuals with less dexterous performance. The anterior ipsilateral PMd enhanced interhemispheric functional coupling consistently with all of the contralateral sensorimotor cortices during the complex task. In contrast, in older group, all of the ipsilateral cortices including the M1 activated during the complex task, however, none of the cortices showed performance-related activity change. Increase in functional connectivity within the contralateral sensorimotor cortices rather than interhemispheric connectivity was observed during the complex task.

The results suggest the importance and complementary role of the ipsilateral PMd for complex finger movement in the younger brains, and the strategic difference (interhemispheric vs. intrahemispheric) when the younger and aging brains perform the movement.

## 1. Introduction

The human sensorimotor system is full of plasticity. For example, when the brain or the spinal cord is injured, the brain often recruits normally-underused cortical sensorimotor areas (dorsal premotor cortex [PMd], primary motor cortex [M1], primary somatosensory cortex [S1], and superior parietal cortex of Area 2) during ipsilateral hand/finger movements (Grefkes & Fink, 2020; Hoy et al., 2004; Jang et al., 2004; Lotze et al., 2006; Ward et al., 2008). Recently, animal studies have suggested that such recruitment of ipsilateral sensorimotor cortices is mediated by disinhibition of interhemispheric inhibition between the left and right motor cortices (Yamaguchi et al., 2023) by acetylcholine modulating GABAergic interneurons (Handa et al., 2024). In stroke patients, jamming transcranial magnetic stimulation (TMS) to the ipsilateral sensorimotor cortices disrupts finger movement, especially the TMS to the PMd disrupts the movement more than when it is applied to the M1 and the superior parietal lobule (Lotze et al., 2006). It is also shown in non-human primates that the ipsilateral PMd is an important node in the recovery of grasping function after the spinal cord injury (Chao et al., 2019). These clinical evidences suggest that the ipsilateral sensorimotor cortices are capable of complementing sensorimotor control of finger movement, especially the ipsilateral PMd plays a particularly important role.

In healthy younger adults, the ipsilateral sensorimotor activation has also been reported when they perform complex motor tasks that require coordination between fingers (Loibl et al., 2011; Uehara et al., 2012; Verstynen et al., 2005). However, it is unclear whether the ipsilateral PMd plays particularly important roles among the ipsilateral sensorimotor cortices by showing complementary activity also when they perform such complex motor task, as shown in the above clinical studies. In the present study, we tested the hypothesis that the ipsilateral PMd becomes active when healthy younger adults perform a complex motor task that requires coordination between fingers, in a way that its activity complements poor motor performance of the task. In addition, it is shown that greater ipsilateral sensorimotor activation is accompanied by greater functional connectivity between the hemispheres within the sensorimotor network when younger adults perform a demanding motor task (Andrushko et al., 2021). So, if the PMd plays particularly important roles when they perform the complex task among the ipsilateral sensorimotor cortices, we may expect that the PMd is a particularly important region which augments interhemispheric functional connectivity with the contralateral sensorimotor cortices during the task.

In contrast, in healthy older adults, the ipsilateral sensorimotor activation can be seen not only during the complex motor task (Loibl et al., 2011) but also during simple unilateral finger movement (Hutchinson et al., 2002; Riecker et al., 2006) and even during a non-motor kinesthetic stimulation to the hand (Naito et al., 2021). Hence, the view that the brain recruits the ipsilateral sensorimotor cortices to complement complex finger motor control is controversial in the case of older adults. If so, even though ipsilateral sensorimotor activation can be seen during complex finger movement in older adults, such activation might not be related to motor complementation in their aging brains. In addition, since it is known that transcallosal fiber (Fling & Seidler, 2012; Lebel et al., 2012; Ota et al., 2006; Strauss et al., 2019; Sullivan et al., 2006) and interhemispheric inhibition (Davidson & Tremblay, 2013; Fling et al., 2011; Naito et al., 2021; Talelli et al., 2008) between the bilateral sensorimotor cortices are deteriorated in older adults, it is unlikely that an aging brain uses the same interhemispheric strategy that a younger brain uses to perform the complex task.

In the current work, using functional magnetic resonance imaging (fMRI), we addressed these questions by measuring brain activity during a complex motor task requiring coordination between the right fingers and during a simple motor task with the right index finger. We first identified brain regions that increased activity during the complex task as compared to the simple task. Since we are particularly interested in the sensorimotor cortices (PMd, M1, S1, and Area 2), we defined each of these regions as region-of-interest (ROI) in each hemisphere, and carefully examined activation and deactivation in each hemisphere. Next, we examined if the ipsilateral sensorimotor activity emerges in relation to poor motor performance during the complex task. Finally, we examined brain regions that enhanced functional coupling with each contralateral seed region (PMd, M1, S1, and Area 2) during the complex task as compared to the simple task. We investigated these points both in younger and older adults to elucidate differences between them.

## 2. Materials & Methods

### 2.1 Participants

Healthy right-handed younger adults (YA group: 22 men, 9 women: mean age, 22.1 ± 1.8) and older adults (OA group: 31 men, 17 women: mean age, 71.1 ± 4.3) participated in this study. We assessed the cognitive status of older participants using the Mini-Mental State Examination (MMSE). All participants scored higher than the cut-off score of 24 (Lopez et al., 2005). We confirmed the handedness of the participants using the Edinburgh Handedness Inventory (Oldfield, 1971). We verified the absence of participants with a history of neurological, psychiatric, and motor disorders based on their self-report.

The study protocol was approved by the Ethics Committee of the National Institute of Information and Communications Technology, and the MRI Safety Committee of the Center for Information and Neural Networks (CiNet; no. 2003260010). We explained the details of the present study to all participants before the experiment, and they then provided written informed consent. The study was conducted according to the principles and guidelines of the Declaration of Helsinki (1975).

### 2.2 Motor tasks

The participants were placed in the supine position in the MRI scanner, and performed two motor tasks (simple and complex motor tasks). Their heads were immobilized using sponge cushions and adhesive tapes, and their ears were plugged. Both the left and right hands were placed on towels and cushions with their palms down so that the arm would not be strained. To ensure that the number of movements matched across participants during the measurement of brain activity, they were asked to perform movements to a constant periodic sound. They were also instructed to close their eyes, relax their entire bodies without producing unnecessary movements and think only of things relevant to the tasks assigned.

1. *Simple task*: The participants were asked to press an MR-compatible button (Current Design Inc., Philadelphia, PA) with their right index finger in synchronization with a 1 Hz sound generated by a computer (Figure 1a left panel). Throughout an fMRI run, the participants kept their right index fingers on the button, and performed repetitive button pressing without releasing the finger from the button. The coefficient of variation (CV) of the button pressing interval was used as a measure of performance for the simple task to evaluate the variability of button pressing (Figure 1b left panel).
2. *Complex task*: The participants were asked to rotate a 9.8 cm, 21 g wooden stick 180° counterclockwise with their right fingers (thumb, index, and middle fingers) in synchronization with a 0.8 Hz sound generated by the computer (Figure 1a right panel). We selected this task because this can be considered as a complex motor task that requires coordination between the fingers (see more in discussion). From preliminary experiments, some older participants had difficulty rotating the stick at 1 Hz. Therefore, we chose a rate of 0.8 Hz, which allowed even these individuals to perform the task successfully. We confirmed that the participants performed 0.8-Hz repetitive rotation of the stick by visual inspection throughout each fMRI run. In addition, each participant’s maximum performance on the complex task was also measured outside the MRI scanner. The participants were required to rotate the stick as many times as possible in 10 seconds, and the number of 180° rotations was counted. Three trials of this task were performed, and the average number of rotations was used as a measure of each participant’s maximal performance (Figure 1b right panel).

**Figure 1.**
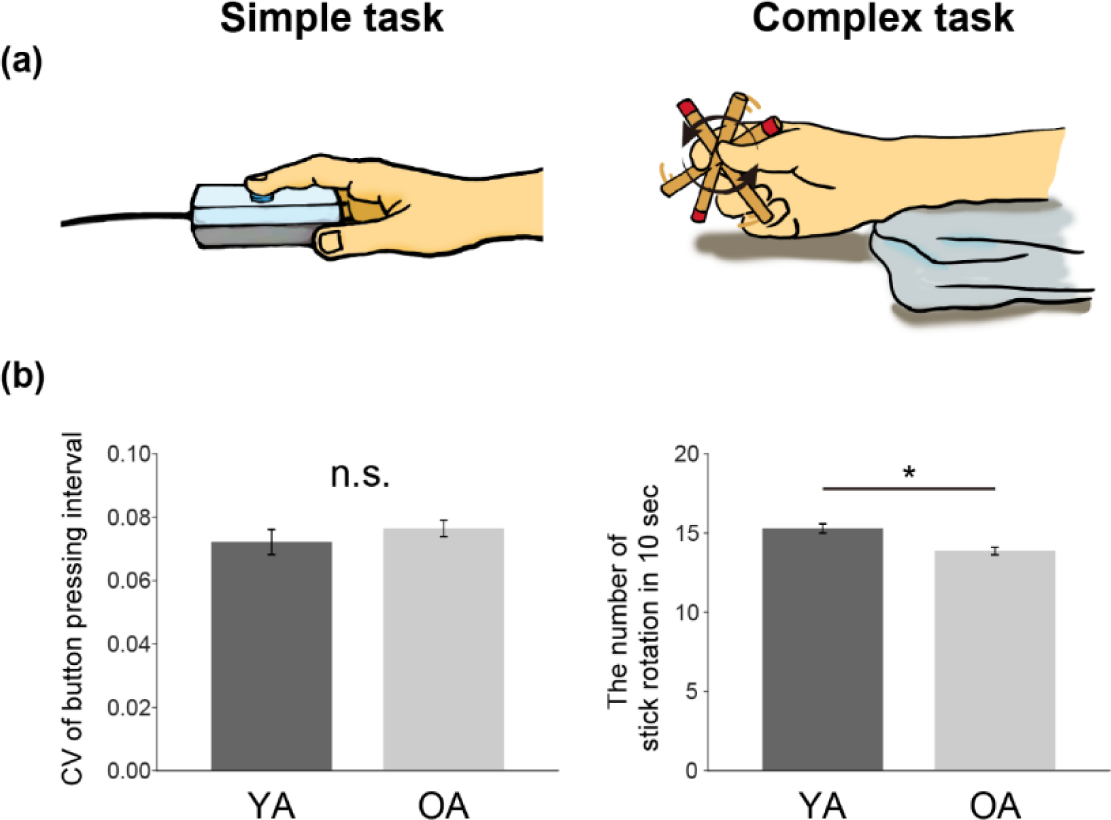
Motor tasks (a) and their performances (b). (a) Simple task: button pressing with the right index finger (left panel). Complex task: stick spinning with the right thumb, index, and middle fingers (right panel). (b) Performances of the simple task (left panel) and the complex task (right panel). The dark gray bars represent the data from the YA group, and light gray bars represent the data from the OA group. Error bars indicate standard errors of the mean across participants (SEM). * indicates p < 0.05. Abbreviations: CV, coefficient of variation; YA, younger adult; OA, older adult; n.s., not significant.

The participants performed each task for two sessions, for a total of four sessions. One session consisted of five task epochs and five rest epochs (baseline state) of 15s, each alternating with the task epochs, starting with the rest epoch. In addition, an extra 10 seconds was provided before the first rest epoch for magnetization stabilization. Thus, one session lasted 160s. During the rest epochs, participants received auditory stimuli at a rate of 1 Hz (simple task) or 0.8 Hz (complex task) but did not move their fingers. We instructed the participants to close their eyes just before starting each session because eye closure duration may affect brain activity (Merabet et al., 2007; Weisser et al., 2005). Half of the participants in each group performed the simple task first, while the other half performed the complex task first. During the fMRI run, the participants are given auditory instructions (“3, 2, 1, start” and “stop”) through an MR-compatible headphone to inform them of the start and finish of a task epoch. These instructions were also generated by a computer. The timings of the 1-Hz sounds and button pressing for each participant were computer recorded.

In the behavioral analysis (Figure 1b), we excluded the data obtained from one younger participant for each task, because his data exceeded ± 2 SD of the mean performance of the YA group. Welch’s t-test without equal-variance assumption was used to evaluate the between-group difference in performance of each task.

### 2.3 fMRI acquisition

Functional images were acquired using T2*-weighted gradient echo-planar imaging (EPI) sequences on a 3.0-Tesla MRI scanner (Trio Tim; Siemens, Germany) equipped with a 32-channel array coil. Each volume consisted of 44 slices (slice thickness, 3.0 mm; inter-slice thickness, 0.5 mm) acquired in ascending order, covering the entire brain. The time interval between successive acquisitions from the same slice was 2,500 ms. An echo time of 30 ms and a flip angle of 80° were used. The field of view was 192 × 192 mm^2^ and the matrix size was 64 × 64 pixels. The voxel dimensions were 3 × 3 × 3.5 mm^3^ in the x-, y-, and z-axes, respectively. We collected 65 volumes for each experimental run. As an anatomical reference, a T1-weighted magnetization-prepared rapid gradient echo (MP-RAGE) image was acquired using the same scanner. The imaging parameters were as follows: TR = 1900 ms, TE = 2.48 ms, FA = 9°, field-of-view (FOV) = 256 × 256 mm^2^, matrix size = 256 × 256 pixels, slice thickness = 1.0 mm, voxel size = 1 × 1 × 1 mm^3^, and 208 contiguous transverse slices.

### 2.4 Functional image analysis

#### 2.4.1 Preprocessing

To eliminate the influence of unsteady magnetization during the tasks, the first 4 volumes (10 seconds) of EPI images in each run were excluded from the analysis. Acquired imaging data were analyzed using SPM 12 (default setting: Wellcome Trust Centre for Neuroimaging, London, UK) running on MATLAB R2017a (MathWorks, Sherborn, MA, USA).

EPI images were realigned to correct for head motion. Time series data of the head position during the fMRI experiment were obtained by a rigid body transformation (linear transformation) using the least squares method for six realign parameters (translation along the x-, y-, and z-axes and the rotational displacements of pitch, raw, and roll). Then, the head movements of the participants were evaluated by the framewise displacement (FD) values based on the six parameters (Power et al., 2012). To inspect FD values through all frames of an entire experimental run, we counted the number of frames that had an FD of over 0.9 mm in each participant according to a previous study (Siegel et al., 2014). The two older participants whose FD values exceeded 0.9 mm in more than 5 % of the total volumes were excluded from the following behavioral and imaging analyses. A nonlinear transformation was also performed to correct for distortions in the functional brain images due to inhomogeneities in the magnetic field caused by the participant’s head movement (unwarp). Next, In each participant, the T1-weighted structural image was co-registered to the mean image of all realigned and unwarped EPI images. The individual co-registered T1-weighted structural image was spatially normalized to the standard stereotactic Montreal Neurological Institute (MNI) space (Evans et al., 1994). Applying the parameter estimated in this process, the individual realigned and unwarped images were normalized to the MNI space with 2-mm isotropic voxel size using the SPM12 normalization algorithm.Finally, the normalized images were filtered using a Gaussian kernel with a full-width at half-maximum (FWHM) of 4 mm along the x-, y-, and z-axes.

#### 2.4.2 Single-subject analysis

After preprocessing, we first explored task-related activations and deactivations in each participant with a general linear model (Friston et al., 1995; Worsley & Friston, 1995). For first-level analysis, a design matrix was prepared for each participant. The design matrix contained a boxcar function for the task epoch that was convolved with a canonical hemodynamic response function (HRF). Six realignment parameters were also included in the design matrix as regressors to correct for residual motion-related noise after the realignment. Contrast images showing activation (task > rest) and deactivation (rest > task) for the simple and complex tasks were created for each participant. In the analysis, global mean scaling was not performed to avoid inducing type I error in the assessment of negative blood oxygenation-level dependent (BOLD) responses (Aguirre et al., 1998).

#### 2.4.3 Regions-of-interest (ROIs)

Since our main interest was the bilateral sensorimotor cortices, we defined ROIs in the hand/finger sections of the bilateral PMd, M1, S1, and Area 2. To define ROIs, we combined the publicly available anatomical maps and a functional image obtained from an independent experiment in which 29 healthy right-handed younger adults performed 60-degree flexion-extension of each of the left and right hands at 1 Hz (Supplementary Information). As for the anatomical maps for M1 (areas 4a and 4p), S1 (areas 3a, 3b, and 1), and Area 2, we used cytoarchitectonic probability maps of the Montreal Neurological Institute (MNI) standard brain in the SPM anatomy Toolbox v3.0 (Amunts et al., 2020; Eickhoff et al., 2005). Since the PMd in the current cytoarchitectonic maps is limited to its medial aspect (areas 6d1, 6d2, and 6d3), we used the precentral map in the Harvard-oxford cortical map (Desikan atlas) for anatomical definition of PMd (Desikan et al., 2006). Hand/finger section of M1, S1, or Area 2 in each hemisphere was determined by depicting overlapping section between the functional map and each cytoarchitectonic map. Similarly, hand/finger section of the PMd in each hemisphere was determined by depicting overlapping section between the functional map and the Desikan’s precentral map, except the M1 and S1 ROIs. The defined PMd ROI seems to be located within the preliminary cytoarchitectonic map of area 6 (Ehrsson et al., 2003). Through this procedure, we eventually defined an ROI for the left or right PMd, M1, S1, or Area 2 (Figure 2a). The total number of the voxels was 996, 469, 393, 308 for the left PMd, M1, S1, or Area 2 ROI, and 896, 312, 555, 295 for the right PMd, M1, S1, or Area 2 ROI. These ROIs were used to identify significant brain activity, and in the definition of seed regions in functional connectivity analysis (see below).

**Figure 2.**
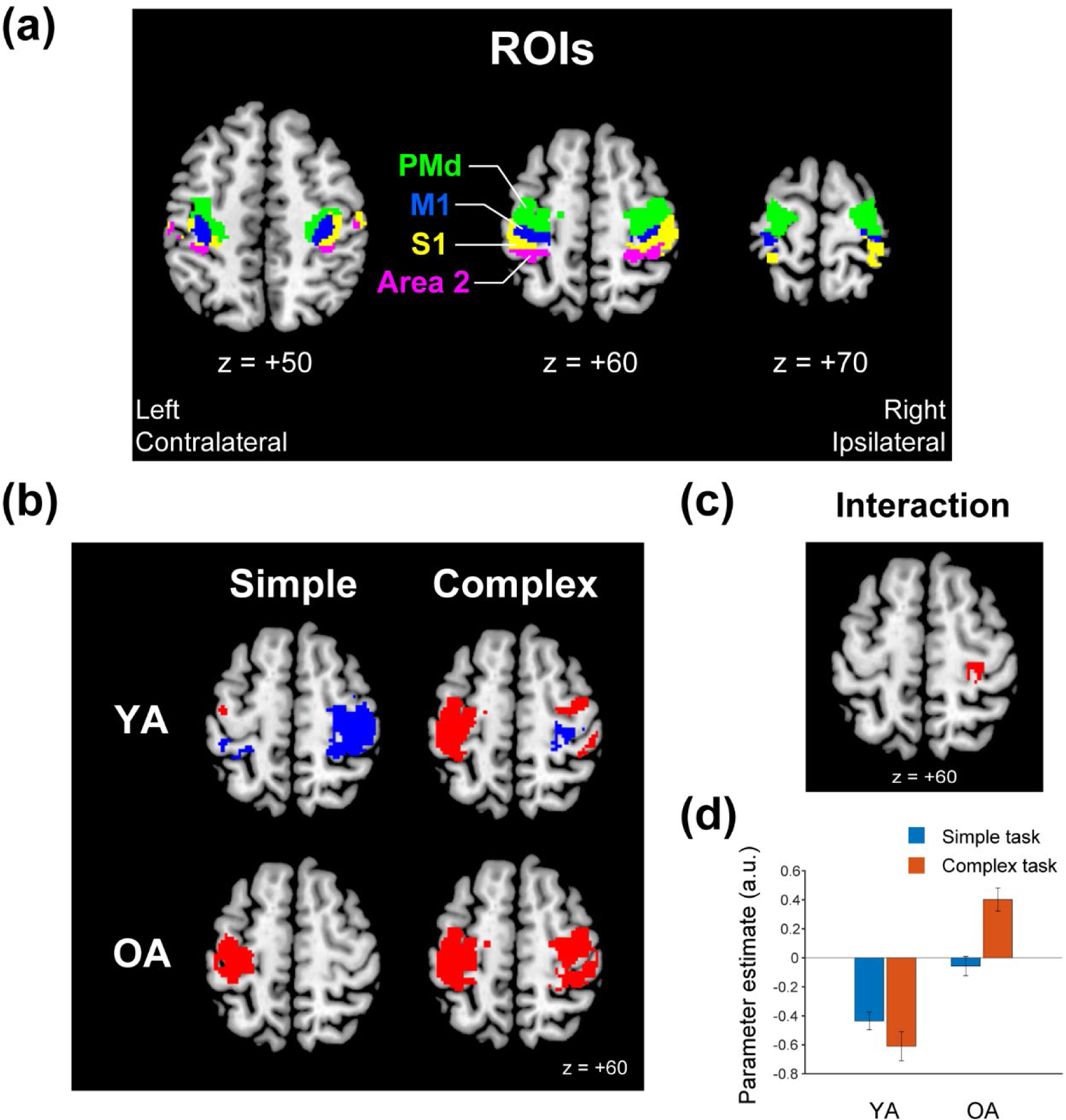
ROIs of the hand/finger sections of the bilateral PMd, M1, S1, and Area 2 (a), and results from the contrast analysis (b-d). (a) Bilateral ROIs [PMd (green), M1 (blue), S1 (yellow), and Area 2 (magenta)] are superimposed on the transverse sections (z = +50, 60, and 70) of the MNI standard brain. (b) Brain activation (red) and deactivation (blue) in the simple (left) and complex (right) tasks in the YA (top row) and OA (bottom row) groups, superimposed on the transverse section (z = +60) of the MNI standard brain. (c) Ipsilateral M1 (red) that showed significant interaction between task and group ([complex task > simple task in OA] > [complex task > simple task in YA]), superimposed on the transverse section (z = +60). (d) Averaged brain activity (parameter estimate) across participants for simple (cyan) and complex (orange) task in the YA and OA groups. Error bars indicate SEM. Abbreviations: MNI, Montreal Neurological Institute; YA, younger adult; OA, older adult; a.u., arbitrary unit; SEM, standard errors of the mean across participants.

#### 2.4.4 Group analysis

For the second-level analysis, we used a full factorial design a within-subject factor (task [2]: complex, simple) and a between-subject factor (group [2]: YA, OA). We first identified activation and deactivation during the simple and complex tasks in each group (Figure 2b). We also investigated differences between tasks (complex > simple) in each group (Supplementary Figure 1). To verify the results, we counted the number of activated and deactivated voxels in each ROI (Supplementary Figure 2). Finally, we examined the interaction between tasks and groups (Figure 2c). In these analyses, we used small volume correction (SVC) approach to identify significant activation and deactivation in the contralateral ROIs composed of the left PMd, M1, S1 and Area 2, and in the ipsilateral ROIs composed of the right PMd, M1, S1 and Area 2, separately. We generated a voxel-cluster image using an uncorrected voxel-wise threshold of p < 0.005, and adopted the family-wise error rate (FWE)-corrected extent threshold of p < 0.05, which was consistently used in the current work. Since we found significant interaction in the right M1 ([complex task > simple task in OA] > [complex task > simple task in YA]), we extracted individual brain activity (parameter estimates) from the significant cluster, and showed the average for each task and group to visualize the interaction effect (Figure 2d). For anatomical identification of the activation and deactivation peaks, we referred to the cytoarchitectonic probability maps (Amunts et al., 2020; Eickhoff et al., 2005).

#### 2.4.5 Correlation analysis

To examine whether the expected ipsilateral sensorimotor activity during the complex task is related to the performance of the complex task, a correlation analysis was conducted using individual maximum performance of the complex task measured outside MRI as a covariate. This was done in each group. In the correlation analysis, one younger participant who was excluded in the above behavioral analysis was also excluded. Since our main interest was the ipsilateral sensorimotor cortices, in the statistical evaluation, we conducted the SVC approach using the ipsilateral ROIs. This analysis revealed significant clusters in the PMd and S1/Area 2 only in the YA group (Figure 3a), Thus, we extracted individual brain activity (parameter estimates) from each cluster, and displayed interparticipant correlation between the brain activity and the maximum performance (Figure 3b). In the case of the OA group, we also displayed the interparticipant correlation using the clusters identified in the YA group (Figure 3c). Finally, we also checked if the contralateral sensorimotor activity during the complex task is related to the maximum performance of the complex task by conducting the SVC approach using the contralateral ROIs (Supplementary Figure 3).

**Figure 3.**
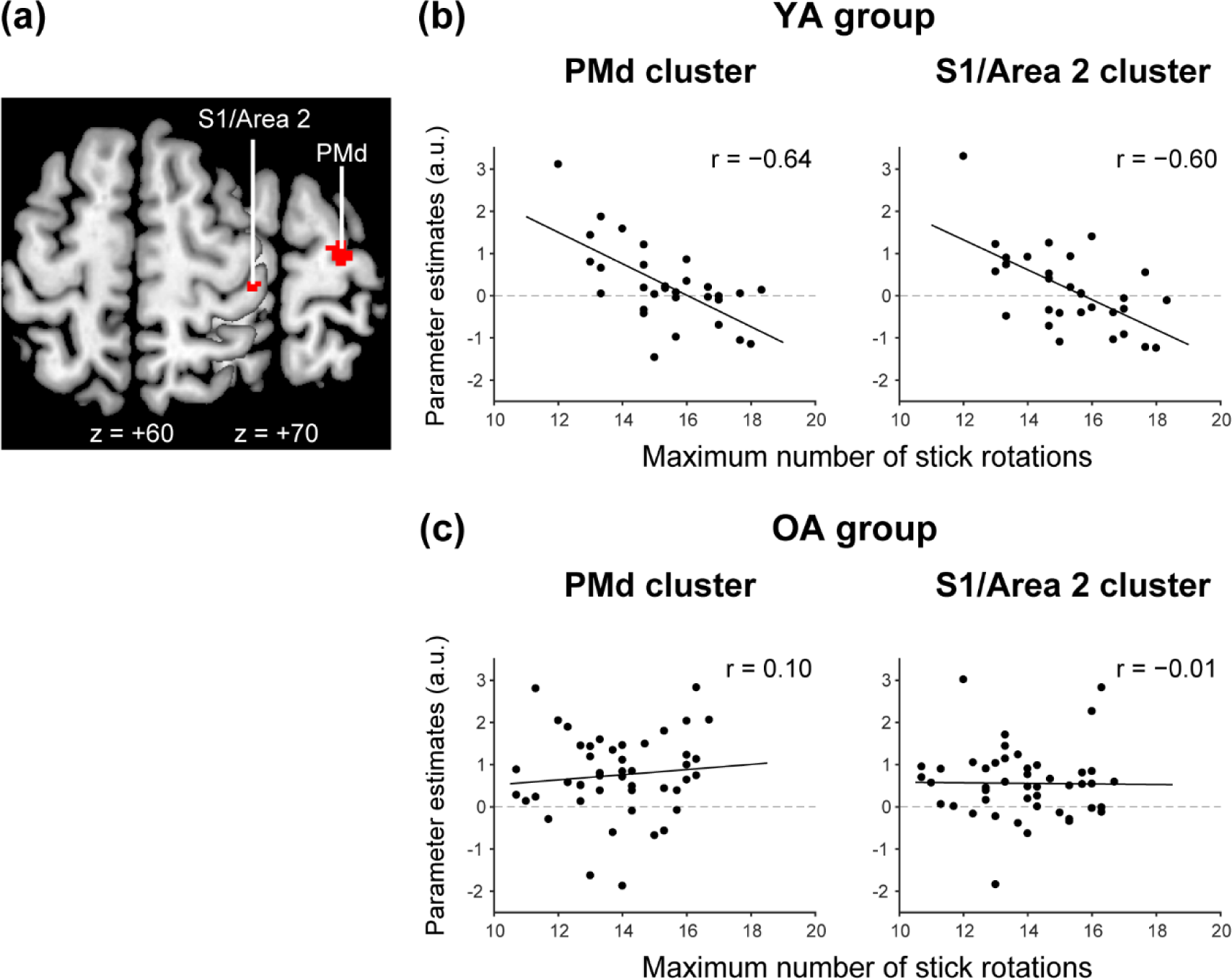
Brain regions in which activity negatively correlated with the performance of the complex task in the YA group (a), and interparticipant correlation between brain activity and the performance in the YA (b) and OA (c) groups. (a) In the YA group, activities in the ipsilateral S1/Area 2 and PMd (red sections) during the 0.8-Hz complex task negatively correlated with the maximum performance of complex motor task measured outside the scanner. These are superimposed on horizontal section of z = +60 and 70 of the MNI standard brain. (b, c) Interparticipant correlation between the performance (x-axis) and brain activity (y-axis) in the YA (b) and OA (c) groups. In b, c, the left panels show correlation between the performance and activity in the PMd cluster, and the right panels show correlation between the performance and activity in the S1/Area 2 cluster. Solid lines in each panel indicate liner regression lines fitted to the data. Abbreviations: MNI, Montreal Neurological Institute; YA, younger adult; OA, older adult; a.u., arbitrary unit.

#### 2.4.6 Task-related functional connectivity analysis

We examined brain regions in which activity enhanced functional coupling with contralateral seed regions (see below) during the complex task as compared to the simple task, by conducting a generalized psychophysiological interaction analysis (gPPI) (McLaren et al., 2012). This analysis was performed on preprocessed fMRI data using the CONN toolbox version 20.b (Nieto-Castanon, 2020; Whitfield-Gabrieli & Nieto-Castanon, 2012). Physiological noises originating from the white matter and cerebrospinal fluid (CSF) were removed using the component-based noise correction method (CompCor) in the toolbox (Behzadi et al., 2007). Head motion-related artifacts, scrubbing, and condition effects were also removed. A temporal band-pass filter of 0.008–0.09 Hz was applied, because we wanted to examine task-related functional connectivity change in this slower range of brain activity fluctuation below than the cardiac and respiratory cycles (0.1–1.2 Hz) (Cordes et al., 2001). We prepared four seed regions in the contralateral PMd, M1, S1, and Area 2. First, we identified brain regions consistently active during the complex task between the YA and OA groups by performing a conjunction analysis (uncorrected voxel-wise threshold of p < 0.005 and extent threshold of p < 0.05 corrected) (Price & Friston, 1997). Each seed region was determined by depicting overlapping region between this functional image and the left PMd, M1, S1 or Area 2 ROI (Figure 4).

**Figure 4.**
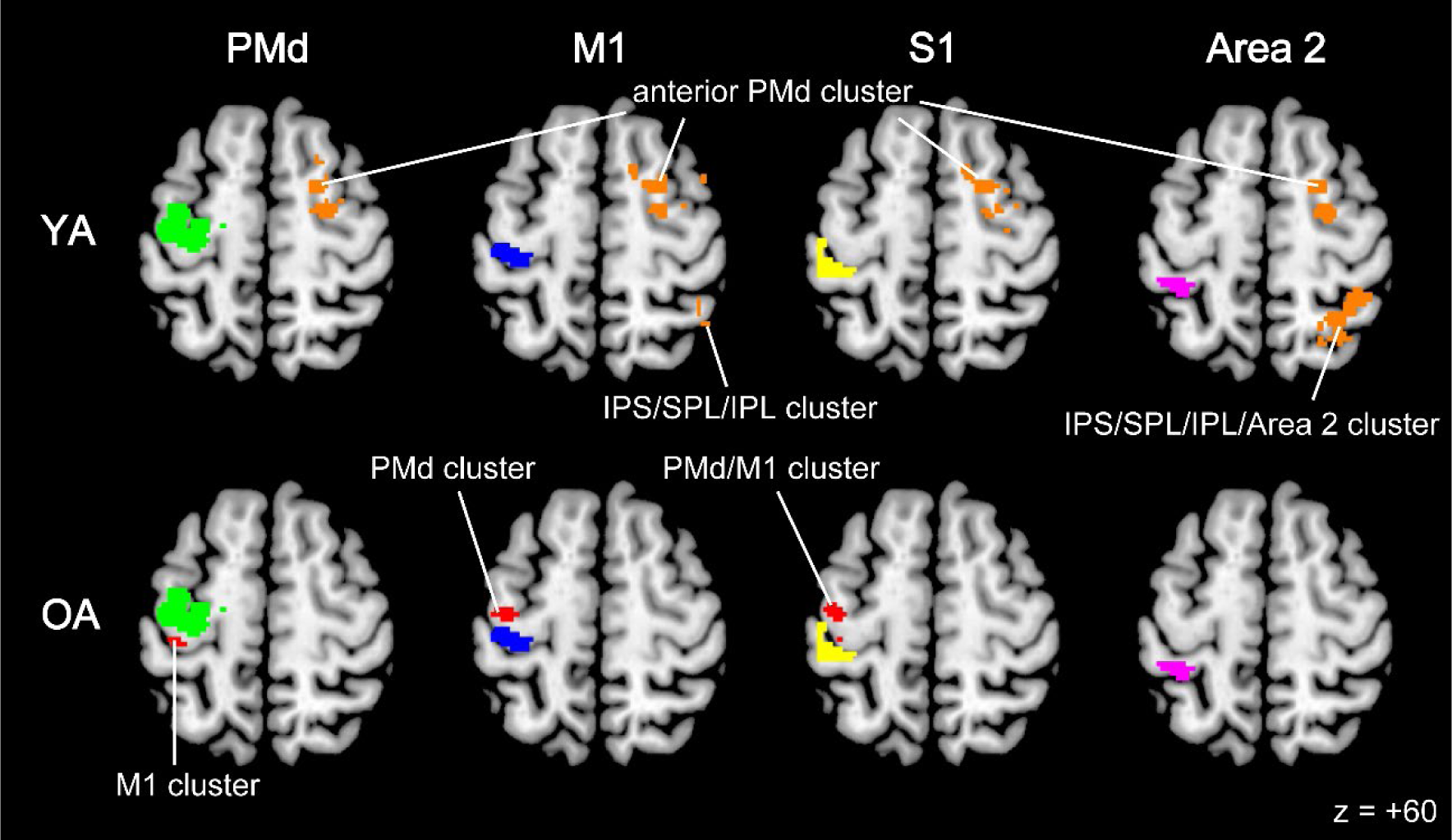
Brain regions in which activity increased functional connectivity with contralateral seed regions during the complex task compared to the simple task in each group. The ipsilateral anterior PMd (orange sections) enhanced functional coupling consistently with all seed regions in the YA group (top row), while brain regions (red sections) in which activity enhanced functional coupling with the PMd, M1, and S1 seeds were observed within the contralateral ROIs in the OA group (bottom row). Green, blue, yellow, and magenta sections represent the contralateral seed regions (PMd, M1, S1, and Area 2), respectively. The activities are superimposed on the transverse section (z = +60) of the MNI standard brain. Abbreviations: MNI, Montreal Neurological Institute; YA, younger adult; OA, older adult; PMd, dorsal premotor cortex; M1, primary motor cortex; IPL, inferior parietal lobule; IPS, intraparietal sulcus area; SPL, superior parietal lobule.

In the gPPI analysis, we used each of the four seed regions. In each participant, the time course of the average fMRI signal across the voxels in each seed region was deconvolved using the canonical HRF (physiological variable). Then, we performed a general linear model analysis using the design matrix and included the following regressors: physiological variable, boxcar function for the task epoch (psychological variable), and multiplication of the physiological variable and the psychological variable (PPI). These variables were convolved with a canonical HRF. Six realignment parameters were also included in the design matrix as regressors of no interest.

In each task, we first generated an image of voxels showing to what extent their activities changed with the PPI regressor of each seed region in each participant. Then, we generated a contrast image (complex task > simple task) that shows complex task-related connectivity change for each participant. We used this individual image in the second-level group analysis, in which the task order was also included as nuisance covariates, to exclude the possibility that the factor affect the results since the order was counterbalanced across participants. In the second-level analysis, in each group, we searched for significant clusters in the contralateral ROIs (the left PMd, M1, S1 and Area 2), and in the ipsilateral ROIs (the right PMd, M1, S1 and Area 2), separately. In the YA group, no significant clusters were identified in the ROIs. However, since we found significant clusters in the entire brain (in motor-related areas just outside the ipsilateral ROIs), we reported these clusters (uncorrected voxel-wise threshold of p < 0.005 and extent threshold of p < 0.05 FWE-corrected) in the YA group (Figure 4).

## 3 Results

### 3.1 Motor performance

In the simple task, both groups successfully performed the 1 Hz button pressing in the scanner. The CV of the button pressing interval, a measure of performance of the simple task, was 0.076 ± 0.031 in the YA group and 0.076 ± 0.018 in the OA group (Figure 1b left panel). Welch’s t-test showed no significant between-group difference (t(54.23) = −0.90, p = 0.37). In the complex task, we visually confirmed that both groups could perform the 0.8 Hz stick rotation in the scanner. However, when we evaluated the maximum number of stick rotation in 10 seconds outside the scanner, this measure of performance of the complex task was 15.20 ± 1.88 times in the YA group and 13.86 ± 1.66 times in the OA group (Figure 1b right panel). Welch’s t-test revealed significantly higher maximum performance in the YA group than in the OA group (t(62.57) = 3.89, p = 2.0 × 10^−4^). Hence, even though the OA group was able to rotate at least 10 times, which would guarantee that the OA group could manage to perform the 0.8 Hz stick rotation in the scanner as did the YA group, their maximum performance was significantly poorer than that of the YA group.

### 3.2 Activation and deactivation in the YA and OA groups

In the YA group, the simple task activated the PMd/M1 and deactivated Area 2 in the contralateral ROIs, and deactivated all areas (PMd, M1, S1, and Area 2) in the ipsilateral ROIs (Figure 2b top left). The complex task activated all areas in the contralateral ROIs. However, within the ipsilateral ROIs, this task activated the PMd, S1, and Area 2, while the M1 remained deactivated as in the simple task (Figure 2b top right). Peaks of activation and deactivation are summarized in Table 1. Eventually, the complex task activated all areas in the contralateral ROIs and the PMd, S1, and Area 2 in the ipsilateral ROIs more strongly than the simple task (Supplementary Figure 1 and Supplementary Table 1). In the OA group, the simple task activated all areas (PMd, M1, S1, and Area 2) in the contralateral ROIs. Unlike the YA group, deactivation in the ipsilateral ROIs disappeared (Figure 2b bottom left). The complex task activated all areas in both contralateral and ipsilateral ROIs (Figure 2b bottom right). Eventually, the complex task activated all areas in the bilateral ROIs more strongly than the simple task (Supplementary Figure 1). These results were validated when we counted activated and deactivated voxels in each ROI (Supplementary Figure 2).

**Table 1.**
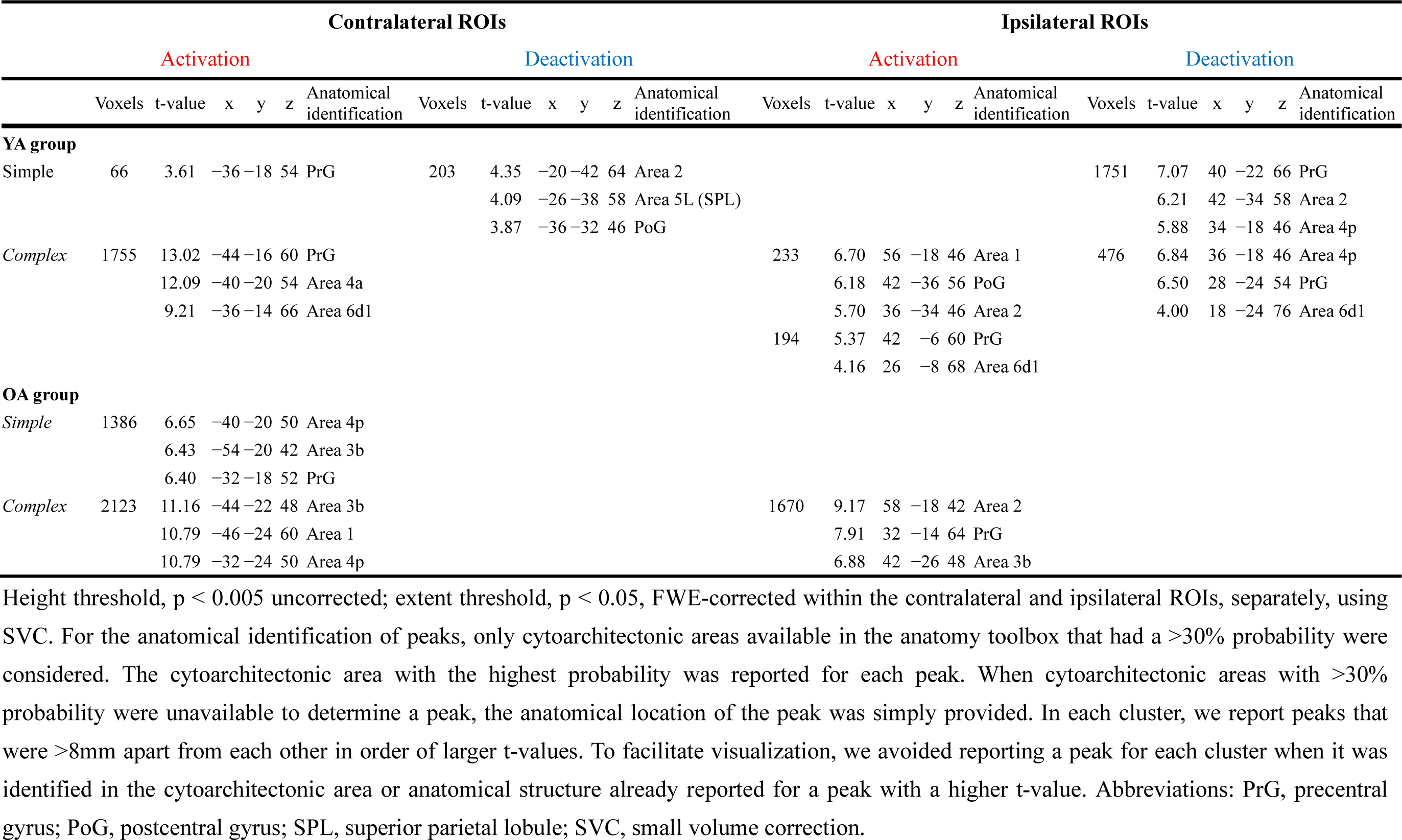
Activation and deactivation during the simple and complex tasks within the bilateral ROIs in each group.

When we examined the interaction ([complex task > simple task in OA] > [complex task > simple task in YA]), we found a significant cluster (223 voxels; peak coordinates = 28, −24, 54, ipsilateral M1) within the ipsilateral ROIs, whereas no significant cluster was found in the contralateral ROIs. This could be due to that, in the OA group, the ipsilateral M1 activated during the complex task, while, in the YA group, this remained deactivated during the complex task as in the simple task (Figure 2d). Activity peaks are summarized in Table 2.

**Table 2.** Areas that showed significant interaction between task and group.

| Size | t-value | x | y | z | Anatomical identification |
| --- | --- | --- | --- | --- | --- |
| 223 | 4.72 | 28 | -24 | 54 | PrG |
|  | 3.72 | 34 | -18 | 48 | Area 4p |
|  | 3.56 | 26 | -32 | 62 | Area 3b |

Height threshold, $p < 0.005$ uncorrected; extent threshold, $p < 0.05$ , FWE-corrected within the contralateral and ipsilateral ROIs, separately, using SVC. For the anatomical identification of peaks, only cytoarchitectonic areas available in the anatomy toolbox that had a $>30\%$ probability were considered. The cytoarchitectonic area with the highest probability was reported for each peak. When cytoarchitectonic areas with $>30\%$ probability were unavailable to determine a peak, the anatomical location of the peak was simply provided. In each cluster, we report peaks that were $>8\text{mm}$ apart from each other in order of larger t-values. To facilitate visualization, we avoided reporting a peak for each cluster when it was identified in the cytoarchitectonic area or anatomical structure already reported for a peak with a higher t-value. Abbreviation: PrG, precentral gyrus; SVC, small volume correction.

### 3.3 Brain regions in which activity correlated with maximum performance of the complex task

We examined brain regions in which activity during the 0.8 Hz complex task correlated with the maximum performance of the complex task outside the scanner, within the ipsilateral ROIs. In the YA group, we found two clusters of voxels in which activity negatively correlated with the performance. One was located in the PMd, and the other was located in the S1/Area 2 (Figure 3a). Peaks of significant clusters are summarized in Table 3. These regions partially overlapped with the regions active during the complex task (Figure 2b top right). When we looked at brain activity, activity in both clusters increased in individuals who showed poorer motor performance outside the scanner (Figure 3b). In contrast, in the OA group, even though all areas in the ipsilateral ROIs including the M1 activated during the complex task (Figure 2b bottom right), none of them showed such significant correlation with the maximum performance (Figure 3c).

**Table 3.** Brain regions in which activity correlated with maximum performance of the complex task.

| Size | t-value | x | y | z | Anatomical identification |
| --- | --- | --- | --- | --- | --- |
| 47 | 4.74 | 24 | -16 | 70 | Area 6d1 |
| 58 | 3.80 | 44 | -22 | 50 | Area 2/Area 3b |
|  | 3.35 | 40 | -18 | 44 | Area 3b |
|  | 3.22 | 44 | -20 | 58 | Area 1 |

Height threshold, $p < 0.005$ uncorrected; extent threshold, $p < 0.05$ , FWE-corrected within the ipsilateral ROIs using SVC. For the anatomical identification of peaks, only cytoarchitectonic areas available in the anatomy toolbox that had a $>30\%$ probability were considered. The cytoarchitectonic area with the highest probability was reported for each peak. When cytoarchitectonic areas with $>30\%$ probability were unavailable to determine a peak, the anatomical location of the peak was simply provided. In each cluster, we report peaks that were $>8\text{mm}$ apart from each other in order of larger t-values. To facilitate visualization, we avoided reporting a peak for each cluster when it was identified in the cytoarchitectonic area or anatomical structure already reported for a peak with a higher t-value. Abbreviation: SVC, small volume correction.

Within the contralateral ROIs, S1/Area 2 activity negatively correlated with the performance in the YA group (Supplementary Figure 3a). The activity increased in individuals who showed poorer motor performance outside the scanner only in the YA group (Supplementary Figure 3b). No regions showed a positive correlation with the performance in either ROI or group.

### 3.4 Enhanced functional connectivity during the complex task in YA and OA group

We examined brain regions in which activity enhanced functional coupling with each seed region (the left PMd, M1, S1, or Area 2) during the complex task compared to the simple task within the contralateral or ipsilateral ROIs. In the YA group, no significant clusters were identified within either ROIs. However, when we searched for clusters in the entire brain, significant clusters just outside the ipsilateral ROIs were identified in motor-related areas (Figure 4). The anterior part of the ipsilateral PMd (partially overlapped with the ipsilateral PMd ROI) enhanced interhemispheric functional coupling consistently with all of the seed regions during the complex task compared to the simple task (Figure 4 top row). Similarly, the ipsilateral intraparietal sulcus area (IPS), superior parietal lobule (SPL), and inferior parietal lobule (IPL) just posterior to Area 2 enhanced interhemispheric functional coupling with the contralateral M1 and Area 2 (Figure 4 top row). Peaks of significant clusters are summarized in Table 4.

**Table 4.**
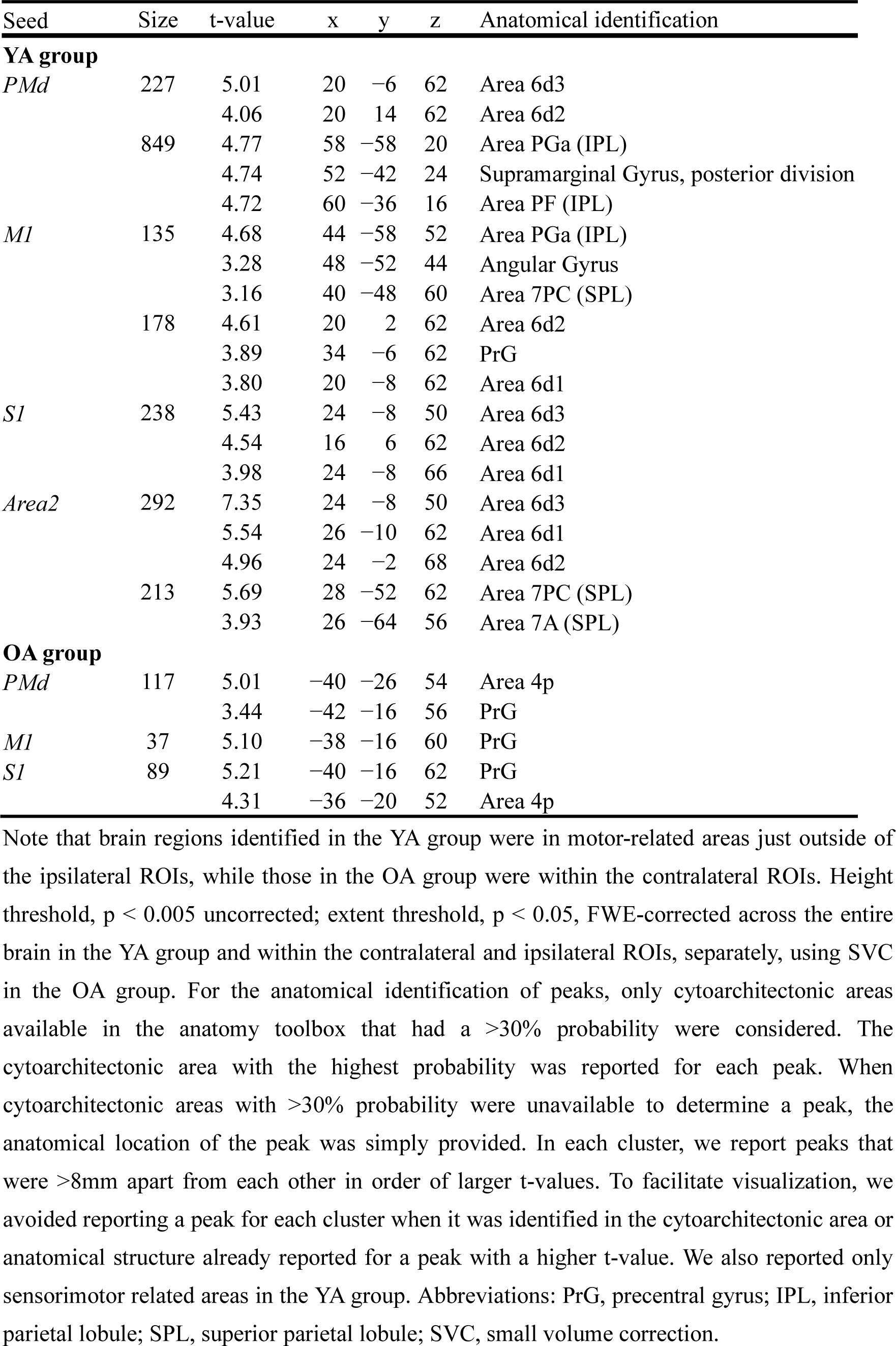
Functional connectivity results (complex > simple)

Unlike the YA group, in the OA group, significant clusters were observed within the contralateral ROIs. The M1 enhanced functional coupling with the PMd during the complex task as compared to the simple task. Similarly, the contralateral PMd enhanced functional coupling with the M1. Finally, the contralateral PMd and M1 enhanced functional coupling with the S1. No significant clusters were identified within the ipsilateral ROIs (Figure 4 bottom row and Table 4) and in the above motor-related areas. Taken all, in the OA group, functional connectivity increased locally within the contralateral hemisphere rather than interhemispheric connectivity.

## 4 Discussion

### 4.1 Younger adults

During the simple task, the ipsilateral sensorimotor cortices were widely deactivated (Figure 2b top left). Ipsilateral sensorimotor deactivation during simple sensory and motor tasks has been repeatedly reported in many previous studies (Allison et al., 2000; Hayashi et al., 2008; Marchand et al., 2007; Morita et al., 2019, 2021; Mullinger et al., 2014; Naito et al., 2021; Newton et al., 2005; Stefanovic et al., 2004), and is thought to be caused by the interhemispheric inhibition from the contralateral cortices (Ferbert et al., 1992; Kobayashi et al., 2003; Mullinger et al., 2014; Talelli et al., 2008).

During the complex task, the ipsilateral sensorimotor cortices were activated (Figure 2b top right), as reported in previous studies which investigated neural correlates of complex and demanding motor tasks (Andrushko et al., 2021; Barany et al., 2020; Buetefisch et al., 2014; Hutchinson et al., 2002; Loibl et al., 2011; Uehara et al., 2012). However, these were observed mainly in the PMd, S1, and Area 2, while the ipsilateral M1 remained deactivated as it was during the simple task. Hence, the current work elucidated clear regional difference in the ipsilateral sensorimotor activation during the complex motor task.

In the current work, we used relatively smaller (4 mm FWHM) Gaussian filter to spatially smooth the functional images. However, this is not a main reason for the regional difference because the same pattern of ipsilateral activation and deactivation was also observed when using larger filer (8 mm) often used in the previous studies (Supplementary Figure 4). In addition, when we further examined brain activity while another nine healthy right-handed younger adults performed the same complex task, we confirmed that all of them consistently showed ipsilateral M1 deactivation, and seven of nine participants showed ipsilateral PMd, and five of nine showed ipsilateral S1/Area 2 activations. As group effect, ipsilateral PMd activation and M1 deactivation were observed, which basically replicates the current results (Supplementary Figure 5).

#### 4.1.1 M1

We adopted the current complex task because this task requires complex stick spinning with fingertips that is an important component when performing a Peg test: a widely-used measure of finger dexterity. Previous studies have shown poor performance of the Peg test in children with immature ipsilateral M1 inhibition and also in older adults with deteriorated ipsilateral M1 inhibition, when compared to younger adults with mature ipsilateral M1 inhibition (Naito et al., 2020, 2021). Hence, ipsilateral M1 inhibition is an important factor for higher finger dexterity. As supporting this prediction, M1 deactivation was observed during the complex task in the YA group who showed relatively better performance (Figure 1b right panel). However, we found no correlation between the degree of M1 deactivation and the performance across participants, which was compatible with our previous report (Naito et al., 2020). Once the ipsilateral M1 inhibition becomes mature, the degree of M1 deactivation does not seem to correlate with the performance.

Though the ipsilateral M1 was deactivated during the current complex task, ipsilateral M1 activation is sometimes reported during other types of demanding hand motor tasks, ex. high force unimanual handgrip (Andrushko et al., 2021; Barany et al., 2020; Buetefisch et al., 2014). Hence, the recruitment of the ipsilateral M1 may be task-dependent, though the ipsilateral M1 deactivation seems to be reproducible as far as using the current complex task (Supplementary Figure 5). In the brain of typically-developed younger adults, there are interhemispheric excitatory and inhibitory circuits between the left and right M1s (Ni et al., 2020). The series of findings suggest that the younger brains can adaptively control movement by flexibly and plastically changing the balance of the interhemispheric facilitation and inhibition between the two M1s.

#### 4.1.2 PMd

In the precentral motor regions, while the ipsilateral M1 was deactivated, the ipsilateral PMd was activated during the complex task (Figure 2b top right). Importantly, as we hypothesized, the participants with lower maximum performance of the complex task recruited more the ipsilateral PMd activity (Figure 3b). One may assume that the 0.8 Hz stick rotation could be more demanding for these participants, and they recruit the activity to complement their lower performance.

As described in Introduction, complementary role of the ipsilateral PMd for the control of hand movement has been shown in the brains of stroke patients (Lotze et al., 2006) and after the spinal cord injury in non-human primates (Chao et al., 2019). In addition, the corticospinal projection from the ipsilateral PMd is known in primates (Kuypers & Brinkman, 1970; Morecraft et al., 2019). Although we should clarify the causal relationship between the ipsilateral PMd activity and the stick rotation performance in future, above lines of evidence imply the possibility that the ipsilateral PMd in the brains of healthy younger adults also capable of complement complex motor control. If this view is correct, ipsilateral PMd recruitment could be a common strategy for the brains to compensate and complement hand motor function not only after spinal cord injury and brain stroke but when healthy younger brains perform complex and demanding hand movements.

In the current work, the importance of ipsilateral PMd during the complex task is also corroborated by the finding that the ipsilateral PMd (anterior part) enhanced functional coupling consistently with all of the contralateral seed regions during the complex task (Figure 4 top raw). This suggests, among the ipsilateral sensorimotor cortices, the PMd is a particularly important region when the contralateral sensorimotor cortices try to communicate with the ipsilateral hemisphere during the complex task. The cluster was mainly located anterior to the region active during the complex task (Figure 2b top right) and to the region in which activity correlated with the performance (Figure 3a). The anterior PMd region seems to well correspond to the region involved in higher-order motor planning/preparation, in concert with the SPL (Furuta et al., 2023; Gerardin et al., 2000; Hanakawa et al., 2008; Solodkin et al., 2004; Stephan et al., 1995). Thus, this anterior region may play slightly different roles from the regions that increased activity during the complex task (Figures 2b and 3a). Anyhow, the series of results support the hypothesis that the PMd plays particularly important roles among the ipsilateral sensorimotor cortices when the younger adult brains perform complex finger movement.

#### 4.1.3 S1 and Area 2

The ipsilateral S1 and Area 2 were also activated during the complex task (Figure 2b top right). The current complex task was stick rotation with three fingers. Thus, it is likely that the brain receives more somatosensory input from the hand/finger muscles and skin when compared to the simple task (button pressing with the index finger). In primates, it is shown that area 2 neurons are characterized by their involvement in the processing of somatosensory information from the bilateral hands (Iwamura et al., 1994), and that human area 2 in each hemisphere responds to proprioceptive stimulation to both hands (Naito et al., 2005). Hence, the present ipsilateral S1 and Area 2 activation together with their contralateral activation (Figure 2b top right) during the complex task might be involved in such complex somatosensory information processing. To support this view, the ipsilateral (Figure 3b) and contralateral (Supplementary Figure 3) S1 and Area 2 activity was negatively correlated with maximum performance of the complex task, meaning that the activity increased in the clumsy participants with lower maximum performance of the complex task. If the 0.8 Hz stick rotation could be more demanding for these participants, we may assume miscellaneous somatosensory input derived from redundant movements due to clumsy control of stick rotation might increase these activities. However, since jamming TMS to the SPL (Area 2) may disrupt finger movement (Lotze et al., 2006), one should bear in mind the possibility that the somatosensory activity somehow contributes to motor control, ex sensory guidance (Rothwell et al., 1982) and/or sensory prediction (Christensen et al., 2007).

#### 4.1.4 Causality analysis using Linear Non-Gaussian Acyclic Model (LiNGAM)

In the YA group, the complex task activated the contralateral sensorimotor cortices and the ipsilateral PMd, S1 and Area 2, while the ipsilateral M1 remained deactivated (Figure 2b top right). In addition, the ipsilateral PMd (especially anterior part) enhanced functional coupling consistently with all of the contralateral seed regions during the complex task (Figure 4). However, these analyses can not provide information about causal relationship between the sensorimotor activities. Therefore, we performed a causality analysis using the Linear Non-Gaussian Acyclic Model (LiNGAM) to explore causal relationship between brain activities across the eight bilateral ROIs (left or right PMd, M1, S1, or Area 2) during the complex task in the YA group (Supplementary Information). Advantage of this approach is that the LiNGAM allows us to explore causal relationship (both positive and negative) between brain activities across multiple brain regions without requiring prior knowledge or specific hypothesis for the network structure (Ogawa et al., 2022). Its drawback is that not all causal relationships obtained from this analysis can be clearly interpreted based on neuroscientific knowledge known to date.

The results are shown in Supplementary Figure 7. Although the right PMd enhanced functional coupling with all of the contralateral sensorimotor cortices (Figure 4), it received positive influences from the contralateral PMd and M1, with stronger influence from the former (Supplementary Figure 7). Since such interhemispheric PMd-PMd interaction plays very important roles when the brain compensates for the damaged contralateral motor pathway during the recovery phase of grasping after the unilateral spinal cord injury in non-human primates (Chao et al., 2019), the present result suggests that the interhemispheric PMd-PMd interaction also plays important roles even when the healthy younger brains complement control of complex finger movement (Figure 3a, b).

The right M1, which was suppressed during the complex task (Figure 2b top right), was ranked lower in causal order among all eight ROIs (Supplementary Figure 7). The right M1 received positive influences from the right PMd and negative (inhibitory) influence from the contralateral side (Area 2). Although several previous studies have reported ipsilateral PMd and M1 activations during complex finger movements (Loibl et al., 2011; Uehara et al., 2012; Verstynen et al., 2005), there seems to be a hierarchical order in their recruitment during complex finger movement: the PMd is recruited relatively immediately, but whether the M1 is recruited or not appears to be determined by the interaction between the positive influence from the ipsilateral PMd and the negative influence from the contralateral sensorimotor cortices.

### 4.2 Older adults

In the OA group, the broader ipsilateral sensorimotor deactivation in the YA group disappeared during the simple task (Figure 2b bottom left), which fits well to previous reports (Hutchinson et al., 2002; Mattay et al., 2002; Morita et al., 2021; Naito et al., 2021; Riecker et al., 2006; Talelli et al., 2008). Aging-related reduction/loss of ipsilateral sensorimotor deactivation is also reported during non-motor kinesthetic stimulation of the unilateral hand (Naito et al., 2021). Thus, reduction/loss of ipsilateral sensorimotor deactivation can occur independent of motor control, and may simply reflect aging-related reduction of interhemispheric inhibition. Exact neural mechanisms underlying this phenomenon are unveiled. However, if one may assume that the ipsilateral sensorimotor deactivation is associated with local neural inhibition mediated by inhibitory neurotransmitter (GABA), aging-related reduction of GABA concentration (Gao et al., 2013) could be an important factor for the aging-related reduction/loss of ipsilateral sensorimotor deactivation.

Unlike the YA group, the complex task activated all areas in the ipsilateral ROIs including the ipsilateral M1. Nevertheless, none of these areas showed a correlation between the individual activity level and performance (Figure 3c). Thus, activity increase in the ipsilateral sensorimotor cortices in the OA group has nothing to do with the performance of the complex task and with complementation of their clumsy finger movements (Figure 3c). Hence, increase in the ipsilateral M1 activity during the complex task in the OA group (Figure 2d) could merely an epiphenomenon resulting from their reduced or lost inhibition from the left M1 to the right M1. The reduction/loss of this interhemispheric inhibition in the present older participants seems to be supported by the fact that involuntary movements of the left finger were observed in several older adults while performing the complex task in the scanner, which can be called mirror movement (Carson, 2005) or mirror overflow (Luo et al., 2022) due to weakening of interhemispheric inhibition (Hoy et al., 2004).

It is well-established that transcallosal fiber between the left and right M1s is quantitatively and qualitatively degraded in older adults (Fling & Seidler, 2012; Lebel et al., 2012; Ota et al., 2006; Strauss et al., 2019; Sullivan et al., 2006). Thus, recruiting interhemispheric regions to perform a complex motor task is not a good strategy for the aging brains with degraded interhemispheric fiber. Instead, it seems that the aging brains tend to take the strategy to increase short-range functional connectivity within the contralateral sensorimotor cortices. This can be said an infantilized strategy, because functional brain networks generally develop from local (short-range) to distant (long-range) organization (Amemiya et al., 2019; Dosenbach, 2010; Fair et al., 2009). Furthermore, the stroke brains often take the strategy to recruit ipsilateral (= contralesional) sensorimotor cortices to compensate hand motor function (Grefkes & Fink, 2020). However, if our view is correct, the aging brains could be poor at taking the interhemispheric strategy due to degraded interhemispheric fiber.

A phenomenon called hemispheric asymmetry reduction in older adults (HAROLD) is known in the aging brain (Cabeza, 2002; Cabeza et al., 2018). This is a phenomenon in which the left and right hemispheric lateralization of brain function normally seen in younger adults disappears with aging. The HAROLD in the prefrontal cortex is thought to contribute to the maintenance of cognitive function. For example, it is shown that older adults who recruit the bilateral prefrontal cortex in memory recall tasks perform better than those who use only one hemisphere (Cabeza, 2002; Cabeza et al., 2018), suggesting that the brain may be maintaining higher cognitive function by recruiting bilateral neural resources. The ipsilateral M1 activation during the complex task (Figure 2b bottom right) can be interpreted as a HAROLD phenomenon in a broad sense. However, the ipsilateral M1 activity had nothing to do with the performance of the complex task. Previous studies have shown that (1) ipsilateral sensorimotor activation remained constant even when older adults increase movement frequency of the right index finger (Riecker et al., 2006), (2) ipsilateral sensorimotor activation is associated with longer reaction time (Langan et al., 2010), and (3) reduction/loss of ipsilateral M1 inhibition is associated with lower hand dexterity (Naito et al., 2021). These lines of reports suggest that an aging-related ipsilateral sensorimotor activation (probably due to reduced interhemispheric inhibition) does not benefit task performance, and is rather a sign of lower motor performance. Taken all together, ipsilateral sensorimotor activation does not necessarily complement a motor task in older adults.

## 5 Conclusion

We measured brain activity with fMRI while healthy right-handed younger and older adults performed simple (button pressing with the right index finger) and complex (stick rotation requiring coordination between the right fingers) motor tasks. When the younger adults performed the complex task, the brain utilized interhemispheric strategy of recruiting the ipsilateral PMd, probably to complement the complex motor control. Even though ipsilateral sensorimotor activations were seen in older adults, the aging brain could not use the interhemispheric strategy, but intrahemispheric strategy instead. The current work has advanced comprehensive understating about importance of ipsilateral PMd for complex finger movement in the younger brains with mature interhemispheric inhibition, and about strategic difference (interhemispheric vs. intrahemispheric) when the younger brains and the aging brains with deteriorated interhemispheric inhibition perform the movement.

## Author Contributions

All authors contributed to conceptualization, validation and writing - review editing, and investigation. GM, TM, and JP: formal analysis, TM, JP, and EN: funding acquisition. GM, YT, JP, and EN: visualization. GM, JP, and EN: writing - original draft preparation. TM and EN: project administration, and supervision. All authors contributed to the article and approved the submitted version.

## Acknowledgments

The funding sources were not involved in the study design; collection, analysis, interpretation of data; writing of the report; or the decision to submit the article for publication. The authors are grateful to Dr Tsuyoshi Ikegami for his valuable comments on this work. They also thank Dr Tadashi Isa and Dr Hidenori Aizawa for intensive discussion about neural substrates of disinhibition of interhemispheric inhibition through “Hyper-adaptation” project (JP19H05723), CiNet MRI staff and Mr Susumu Minamiyama for their technical support, and Ms Keiko Ueyama for the illustration.

## Data Availability Statement

Datasets are available on request: The raw data supporting the conclusions of this article will be made available by the corresponding author (EN).

