## Supplementary information for "Younger adult brain utilizes interhemispheric strategy of recruiting ipsilateral dorsal premotor cortex for complex finger movement, but not aging brain"

### 1 Generation of functional images of the hand/finger sections in the bilateral sensorimotor cortices to define ROIs

We collected functional images when an independent group of 29 healthy right-handed younger adults (19 men, mean age 25.4 ± 8.2) continuously exerted 60-degree cyclic extension–flexion movements of each of the left and right hand in synchronization with 1-Hz cyclic tones. We prepared a device to control the range of wrist motion, which was used in our previous study (Morita et al., 2023). A movable hand-rest was mounted on the device, and the hand was fixed on the hand-rest that indicated the wrist angle. Two stoppers were fixed onto the device to control the range of the wrist motion across the task epochs and participants. They were positioned to prevent the wrist from extending beyond the straight (0°) position and flexing beyond 60°. The participants had to touch one of the stoppers (0° or 60°) alternately with the hand-rest in synchronization with the 1-Hz audio tones while making controlled and continuous wrist extension–flexion movements. The left and right hand tasks were performed in a different experimental session.

Functional images were acquired using T2*-weighted gradient echo-planar imaging (EPI) sequences on a 3.0-Tesla MRI scanner (Trio Tim; Siemens, Germany) equipped with a 32-channel array coil. Each volume consisted of 44 slices (slice thickness, 3.0 mm; inter-slice thickness, 0.5 mm) acquired in ascending order, covering the entire brain. The time interval between successive acquisitions from the same slice was 2,500 ms. An echo time of 30 ms and a flip angle of 80° were used. The field of view was 192 × 192 $mm^2$ and the matrix size was 64 × 64 pixels. The voxel dimensions were 3 × 3 × 3.5 $mm^3$ in the x-, y-, and z-axes, respectively. We collected 65 volumes for each experimental run. As an anatomical reference, a T1-weighted magnetization-prepared rapid gradient echo (MP-RAGE) image was acquired using the same scanner. The imaging parameters were as follows: TR = 1900 ms, TE = 2.48 ms, FA = 9°, field-of-view (FOV) = 256 × 256 $mm^2$, matrix size = 256 × 256 pixels, slice thickness = 1.0 mm, voxel size = 1 × 1 × 1 $mm^3$, and 208 contiguous transverse slices.

When performing a task, the participants were asked to close their eyes, relax their entire body, refrain from producing unnecessary movements, and only think of the assigned task. Each participant completed one experimental 160-s run for each task. The run comprised five task epochs, each lasting 15 s. Considering each epoch, the participants continuously exerted cyclic movements for each task in synchronization with cyclic audio tones. The task epochs were separated by 15-s baseline (rest) periods. Each run also included a 25-s baseline period before the start of the first epoch. During the experimental run, we provided the participants with auditory instructions that indicated the start of a task epoch (three, two, one, start). We also provided a 'stop' instruction generated by a computer to notify the participants of the end of each epoch. The participants heard the same cyclic audio tones; however, they did not generate any movement during the rest periods. All the auditory stimuli were provided through an MRI-compatible headphone. An experimenter who stood beside the scanner bed checked if the participants were performing each task properly by visual inspection throughout the run.

The same image preprocessing was performed (see main text), and the normalized images were filtered using a Gaussian kernel with a full-width at half-maximum of 4 mm along the x-, y-, and z-axes. After the single-subject analysis (see main text), we performed the second-level group analysis. We identified activation during the task (task > rest) using the same extent threshold of $p < 0.05$ FWE corrected for a voxel-cluster image with an uncorrected voxel-wise threshold of $p < 0.005$. This was done for the left and right hand tasks, separately. The analysis gave us significant activations in the hand/finger section of the left sensorimotor cortices during the right hand task, and in the section of the right sensorimotor cortices during the left hand task. We used these cluster images to functionally define the hand/finger sections of the bilateral sensorimotor cortices.

**2 Percentage of activated and deactivated voxels in each ROI**

In the group analysis, we generated a voxel-cluster image with an uncorrected voxel-wise threshold of $p < 0.005$ with no extent threshold. We generated activation (task > rest) and deactivation (rest > task) maps for each task in each group. We simply counted the number of activated and deactivated voxels, and the voxel number was divided by the total number of voxels in each ROI (see main text) to compute percentage. The results are shown in Supplementary Figure 2. The results matched well to those obtained from the contrast analysis (Figure 2b).

**3 Reproducibility of the contrast result during the complex task in the YA group when using a spatial Gaussian filter of 8 mm FWHM**

In order to check if the contrast result during the complex task in the YA group (= the ipsilateral PMd, S1/Area2 activations and the ipsilateral M1 deactivation; Figure 2b) is affected by the size (4mm FWHM) of spatial Gaussian filter, we conducted the same analysis on the data when using a spatial Gaussian filter of 8 mm FWHM. The result was perfectly replicated (Supplementary Figure 4). Hence, the result (= clear regional difference in the ipsilateral sensorimotor activation and deactivation during the complex task) was not affected by the spatial filter size.

**4 Reproducibility of the results of the complex task**

In order to check reproducibility of the current result obtained from the complex task, we measured brain activity while 9 other healthy right-handed younger participants (9 men, mean age 24.1 ± 1.2) performed the same complex task (0.8 Hz stick rotation).

Functional images were acquired using T2*-weighted gradient echo-planar imaging (EPI) sequences on a 3.0-Tesla MRI scanner (Vida; Siemens, Germany) equipped with a 64-channel array head-neck coil. A multiband imaging technique was used (multiband factor, 3; Moeller et al., 2010). Each volume consisted of 51 slices (slice thickness, 3.0 mm with no inter-slice thickness) acquired in an interleaved manner, covering the entire brain. The time interval between successive acquisitions from the same slice was 1,000 ms. An echo time of 30 ms and a flip angle of 60° were used. The field of view was 210 × 210 $mm^2$ and the matrix size was 70 × 70 pixels. The voxel dimensions were 3 × 3 × 3 $mm^3$ in the x-, y-, and z-axes, respectively.

As an anatomical reference, a T1-weighted magnetization-prepared rapid gradient echo (MP-RAGE) image was acquired using the same scanner. The imaging parameters were as follows: TR = 1900ms, TE = 2.48ms, FA = 9°, field-of-view (FOV) = 256 × 256 $mm^2$, matrix size = 256 × 256 pixels, slice thickness = 1.0 mm, voxel size = 1 × 1 × 1 $mm^3$, and 208 contiguous sagittal slices.

Participants performed two sessions consisting of 190 seconds each. One session consisted of six task epochs and six rest epochs (baseline state) of 15 s, each alternating with the task epochs, starting with the rest epoch. In addition, an extra 10 seconds was provided before the first rest epoch for magnetization stabilization.

The same image preprocessing was performed (see main text), and the normalized images were filtered using a Gaussian kernel with a full-width at half-maximum of 4 mm along the x-, y-, and z-axes. The single-subject analysis was performed. In this analysis, we used small volume correction (SVC) approach to identify significant activation and deactivation in the contralateral ROIs composed of the left PMd, M1, S1 and Area 2, and in the ipsilateral ROIs composed of the right PMd, M1, S1 and Area 2, separately. Here, we identified activation (task > rest) and deactivation (rest > task) using an uncorrected voxel-wise threshold of $p < 0.005$, and adopted the false discovery rate (FDR)-corrected extent threshold of $p < 0.05$. We reported activation (task > rest) and deactivation (rest > task) using an uncorrected voxel-wise threshold of $p < 0.005$ and extent threshold of $p < 0.05$ FDR-corrected in the bilateral ROIs.

The results are shown in Supplementary Figure 5. All of the participants consistently showed ipsilateral M1 deactivation, and seven of nine participants showed ipsilateral PMd, and five of nine showed S1/Area 2 activations. As group effect, ipsilateral PMd activation and M1 deactivation were observed, which basically replicates the current results.

**5 Causality analysis using Linear Non-Gaussian Acyclic Model (LiNGAM)**

In the YA group, the complex task activated the contralateral sensorimotor cortices and the ipsilateral PMd, S1 and Area 2, while the ipsilateral M1 remained deactivated (Figure 2b top right). In addition, the ipsilateral PMd (especially anterior part) enhanced functional coupling consistently with all of the contralateral seed regions during the complex task (Figure 4). However, these analyses can not provide information about causal relationship between brain activities. Therefore, we performed a causality analysis using the Linear Non-Gaussian Acyclic Model (LiNGAM) to explore causal relationship between brain activities across the eight bilateral ROIs (left or right PMd, M1, S1, or Area 2) during the complex task in the YA group (Supplementary Information). Advantage of this approach is that the LiNGAM allows us to explore causal relationship (both positive and negative) between brain activities across multiple brain regions without requiring prior knowledge or specific hypothesis for the network structure (Ogawa et al., 2022). Its drawback is that not all causal relationships obtained from this analysis can be clearly interpreted based on neuroscientific knowledge known to date. This approach will enhance our understanding of causal relationship among these cortical activities when younger adults performed the complex task.

The Linear Non-Gaussian Acyclic Model (LiNGAM) (Shimizu et al., 2006) is a statistical method to discover causal relationship in data. It estimates directed causal relationships, which can be either positive or negative, within an unknown causal structure among variables, based on observed data.

The LiNGAM is expressed by following equation:

$$x_i = \sum_{k(j)<k(i)} b_{ij} x_j + e_i \; (i = 1, \ldots n), \qquad (1)$$

where, observed variable $x_i$ is expressed by linear summation of other variables $x_j$ $(j \neq i, j = 1, \ldots n)$ and external influence $e_i$ $(i =, \ldots, n)$. A causal ordering of the variables $x_i$is represented by $k(i)$. The external influence $e_i$ is assumed to be an independent and non-Gaussian distribution with zero mean and non-zero variance. This assumption is crucial for determining the causal order among variables. A connection weight $b_{ij}$ indicates a causal effect from observed variable $x_j$ to $x_i$. The equation 1 can be rewritten as

$$\mathbf{x} = \mathbf{Bx} + \mathbf{e}, \qquad (2)$$

where the $\mathbf{x}$ and $\mathbf{e}$ is vector of $x_i$ and $e_i$, respectively. Since LiNGAM assumes an acyclic graph structure, there exists a causal ordering among the observed variables. Consequently, if the rows and columns are rearranged accordingly, the $n \times n$ matrix $\mathbf{B}$, constituted by $b_{ij}$, will be lower triangular. Since many latent confounders (unobserved brain areas) might influence brain activities, we used ParceLiNGAM (Tashiro et al., 2014) to estimate $b_{ij}$ and causal ordering among brain activities in the present study. The ParceLiNGAM is robust against latent confounders compare to original LiNGAM (Tashiro et al., 2014) We used open-source library for

LiNGAM (Ikeuchi et al., 2023).

#### 5.1 Construction of data

To apply the ParceLiNGAM to small-sample fMRI data, we utilized a method that aggregates the data points across participants (Ogawa et al., 2022; Smith et al., 2011; Xu et al., 2014). After preprocessing the fMRI data in the same way as in the main manuscript, we normalized and extracted time series data from ROIs for each participant. To mitigate the impact of outliers, we used a robust scaling method as follows:

$$x_{\text{scale}} = \frac{x - Q_2}{Q_3 - Q_1}, \quad (3)$$

where, $Q_2$ is the median of time series $x$ and $Q_1$ and $Q_3$ represent first and third quantile of $x$, respectively. After normalization, we collected data points across all ROIs during the complex task to construct a data matrix across all participants (Supplementary Figure 6).

#### 5.2 Statistic of causal relationship

We used the bootstrap method to assess the statistical significance of causal effects in the estimated causal relationships. Based on previous studies (Ogawa et al., 2022; Shimizu et al., 2011; Xu et al., 2014), we constructed 1,000 distinct data matrices for bootstrap for each group. Each matrix consists of randomly selected 25 participants for each group. We then conducted the Wald test to evaluate the statistical significance of the causal effects between ROIs. Moreover, we confirmed that whether external influence $\mathbf{e} = (\mathbf{I} - \mathbf{B})\mathbf{x}$ satisfied the assumption of non-Gaussianity, which is one of the assumptions of LiNGAM, using Kolmogorov-Smirnov test ($p < 0.05$).

#### 5.3 Results and Discussion

Supplementary Figure 7 shows causal relationship (order) among the eight ROIs during the complex task in the YA group, revealed by the ParceLiNGAM. Although we can not fully interpret all causal relationships obtained from this analysis, based on neuroscientific knowledge known to date, part of the results can be interpreted in accordance with current neuroscientific knowledge.

During the complex task, the left (contralateral) M1, the executive locus for motor control of the right hand, exerted a fairly stronger positive influence on the left PMd. The left PMd exerted positive influences on multiple regions including the right (ipsilateral) PMd. Although the right PMd enhanced functional coupling with all of the contralateral sensorimotor cortices (Figure 4), it received positive influences from the contralateral PMd and M1, with stronger influence from the former. Since such interhemispheric PMd-PMd interaction plays very important roles when the brain compensates for the damaged contralateral motor pathway during the recovery phase of grasping after the unilateral spinal cord injury in non-human primates (Chao et al., 2019), the present result suggests that the interhemispheric PMd-PMd interaction also plays important roles even when the healthy younger brains complement control of complex finger movement (Figure 3).

The right M1, which was suppressed during the complex task (Figure 2b top right), was ranked lower in causal order among all eight ROIs. Within the right (ipsilateral) sensorimotor cortices, the M1 received positive influences from the PMd and the S1, with stronger influence from the former. Hence, it seems that the right PMd could affect whether the right M1 is recruited or not during the complex task. In reality, however, the right M1 remains suppressed (Figure 2b top right) most probably due to inhibitory (negative) influences from the contralateral side (in this case, from Area 2).

In sum, although several previous studies have reported ipsilateral PMd and M1 activations during complex finger movements (Loibl et al., 2011; Uehara et al., 2012; Verstynen et al., 2005), there seems to be a hierarchical order in their recruitment during complex finger movement: the PMd is recruited relatively immediately, but whether the M1 is recruited or not appears to be determined by the interaction between the positive influence from the ipsilateral PMd and the negative influence from the contralateral sensorimotor cortices.

Unlike the YA group, the OA group showed no significant causal relationships among the ROIs. We speculate that this could be related to excessive bilateral sensorimotor activities during the complex task in the OA group (Figure 2b bottom right). Recently, we have shown that, using a spiking neural network model, excessive activity caused by weaken inhibitory effect within a region in a network results in disruption of information transmission within the network (Park et al., 2023). We assume that the excessive bilateral sensorimotor activities during the complex task in the OA group (Figure 2b bottom right) might disrupt information transmission in the brain network, so that no clear causal relationships among the sensorimotor cortices were observed.

**Supplementary Figures**

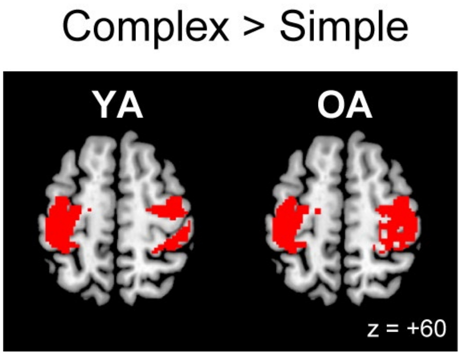


**Supplementary Figure 1. Brain regions more activated during the complex task than the simple task within the contralateral or ipsilateral ROIs in each group**

In the YA group, the complex task activated all areas in the contralateral ROIs and the PMd, S1, and Area 2 in the ipsilateral ROIs more strongly than the simple task. In the OA group, the complex task activated all areas in the bilateral ROIs more strongly than the simple task. Activations are superimposed on the transverse section (z = +60) of the MNI standard brain. Abbreviations: MNI, Montreal Neurological Institute; YA, younger adult; OA, older adult.

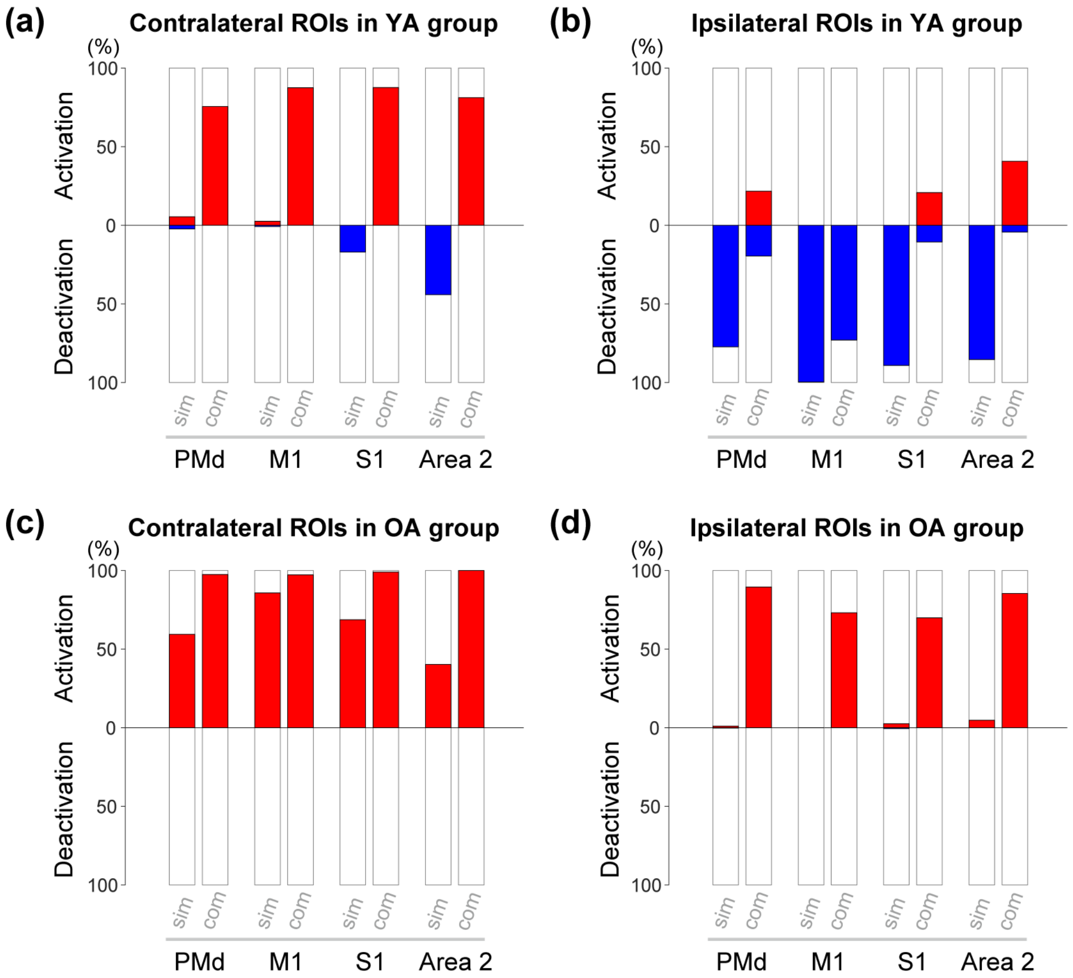


**Supplementary Figure 2. Percentage of activated and deactivated voxels in each ROI.** Results from contralateral (a) and ipsilateral (b) ROIs in YA group. Results from contralateral (c) and ipsilateral (d) ROIs in OA group. Red bars in each panel indicate percentage of activated voxels, and blue bars indicate percentage of deactivated voxels. Overall, the results matched well to those obtained from the contrast analysis (Figure 2b). Abbreviations: YA, younger adult; OA, older adult; sim, simple task; com, complex task.

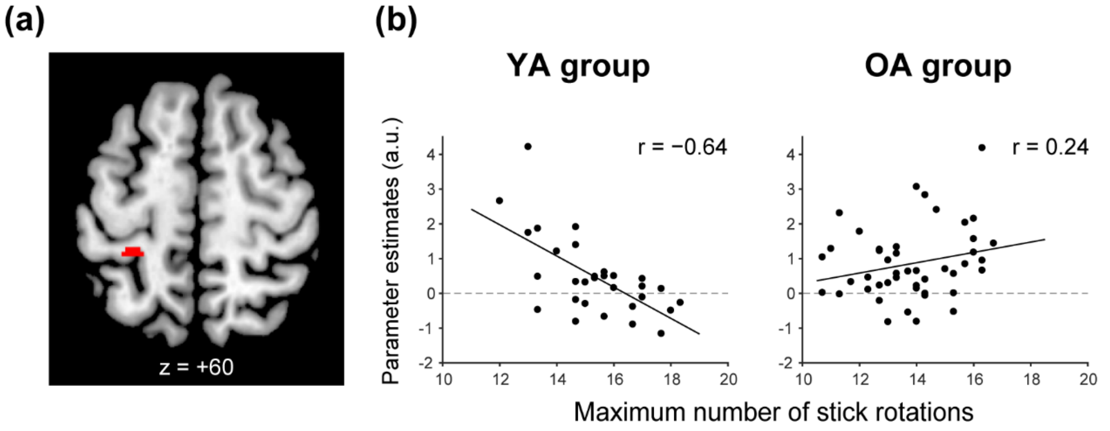


**Supplementary Figure 3. Brain region in which activity negatively correlated with the performance within the contralateral ROIs in the YA group** (a) Activity in the contralateral S1/Area 2 negatively correlated with the performance of complex task in the YA group. The activity was superimposed on horizontal section of z = +60 of the MNI standard brain. (b, c) Interparticipant correlation between the performance (x-axis) and brain activity of the cluster (y-axis) in the YA (b) and OA (c) groups. Solid lines in each panel indicate liner regression lines fitted to the data. Abbreviations: MNI, Montreal Neurological Institute; YA, younger adult; OA, older adult; a.u., arbitrary unit.

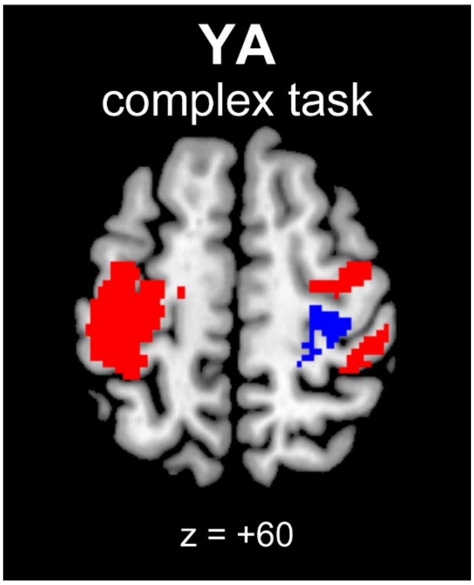


**Supplementary Figure 4. Ipsilateral PMd, S1, Area 2 activation and M1 deactivation when using 8-mm filtered functional images.** Ipsilateral PMd, S1, Area 2 activations (red) and M1 deactivation (blue) were also observed when we used 8-mm filtered images. The results replicated the ipsilateral activation and deactivation pattern observed when analyzing 4-mm filtered images (Figure 2b). Activations and deactivation are superimposed on the horizontal section of z = +60 of the MNI standard brain. Abbreviations: MNI, Montreal Neurological Institute.

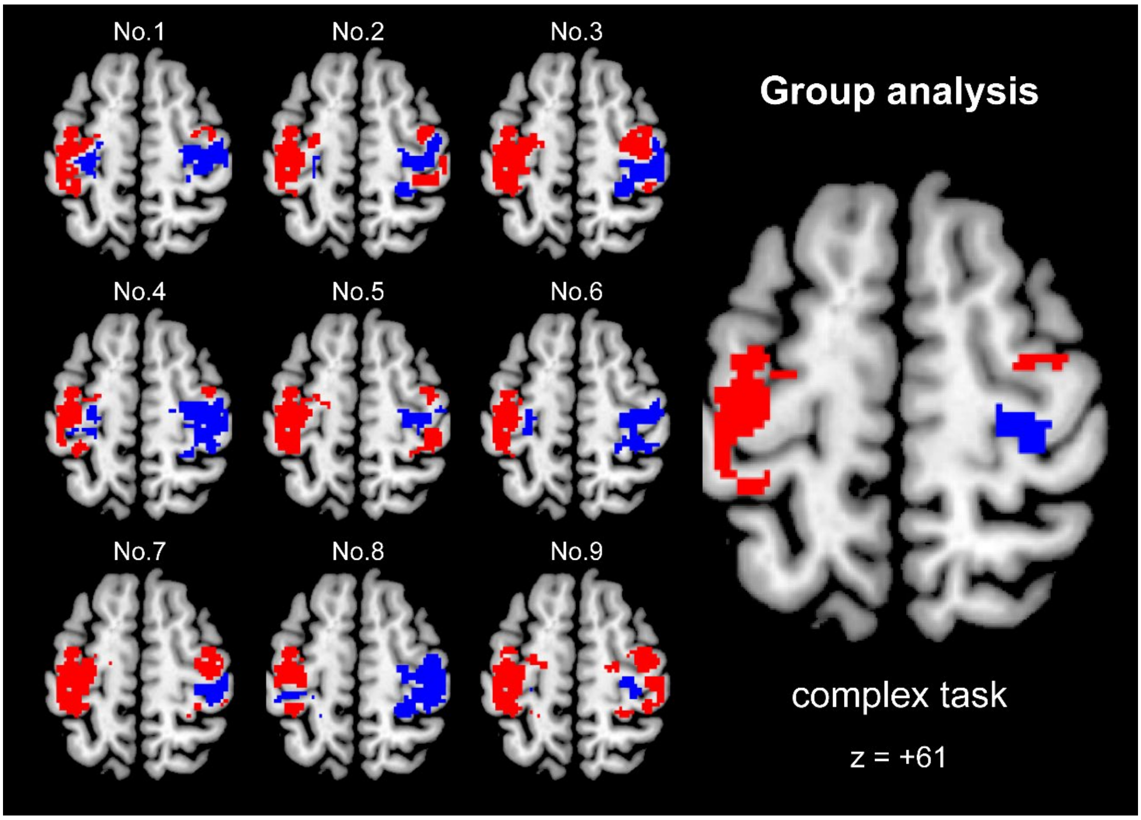


**Supplementary Figure 5. Individual brain activations and deactivations when other nine younger adults performed the complex task, and their group effect.** Activations (red) and deactivations (blue) in individual participants and their group effect are superimposed on horizontal section of z = +61 of the MNI standard brain. Abbreviations: MNI, Montreal Neurological Institute.

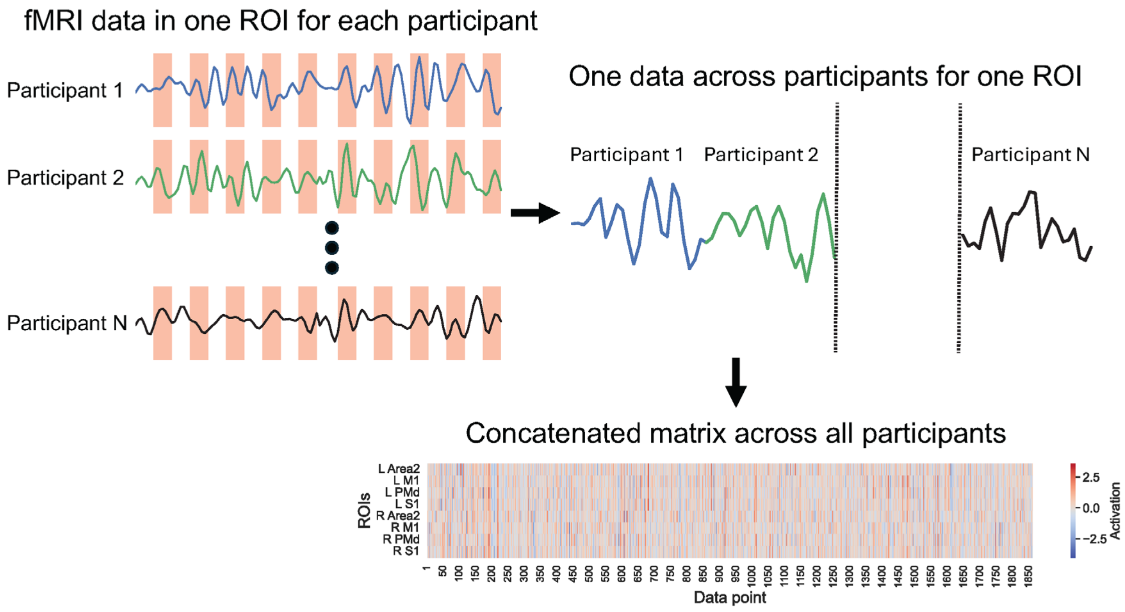


**Supplementary Figure 6. Data matrix using fMRI data.** Areas marked in red indicate the data during the task epoch. The data obtained from each participant were concatenated for each ROI, and this was done for all ROIs to generate the data matrix.

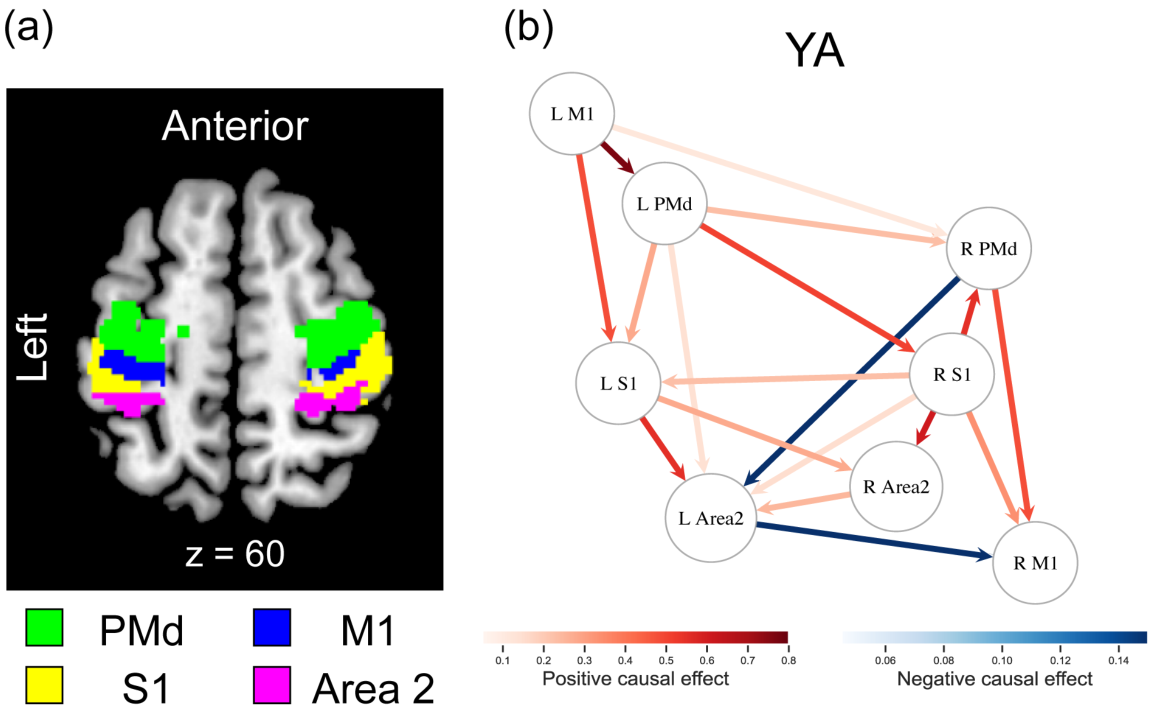


**Supplementary Figure 7. Estimated causal influences (order) among the eight ROIs (a)**

**during the complex task in YA using ParceLiNGAM.** (a) Eight selected ROIs of the left or right PMd, M1, S1 or Area 2 superimposed on horizontal section of z = +60 of the MNI standard brain (see also Figure 2a). (b) Significant causal relationship (order) among the ROIs. Each circle represents a selected ROI. Red and blue arrows indicate the positive and negative causal effects from a ROI to another, respectively. Causal effects are shown only for $p < 0.05$ through the Wald test. Abbreviations: MNI, Montreal Neurological Institute.

**Supplementary Table 1. Brain regions activated more during the complex task than the simple task within the contralateral or ipsilateral ROIs in each group**

| | Size | t-value | x | y | z | Anatomical identification |
|---|---|---|---|---|---|---|
| **YA group** | 1764 | 10.07 | −44 | −18 | 56 | Area 3b |
| | | 9.01 | −30 | −18 | 64 | Area 6d1 |
| | | 8.64 | −32 | −8 | 66 | PrG |
| | 518 | 8.97 | 42 | −34 | 56 | Area 2 |
| | | 7.82 | 52 | −24 | 54 | Area 1 |
| | 453 | 7.54 | 34 | −6 | 64 | PrG |
| | | 6.22 | 26 | −12 | 60 | Area 6d1 |
| **OA group** | 1583 | 8.07 | −46 | −24 | 60 | Area 1 |
| | | 7.27 | −38 | −30 | 50 | Area 4p |
| | | 7.25 | −44 | −34 | 58 | Area 3b |
| | 1704 | 7.59 | 30 | −12 | 58 | Area 6d1 |
| | | 7.25 | 36 | −14 | 66 | PrG |
| | | 7.21 | 34 | −34 | 46 | Area 2 |

Height threshold, $p < 0.005$ uncorrected; extent threshold, $p < 0.05$, FWE-corrected within the contralateral and ipsilateral ROIs, separately, using SVC. For the anatomical identification of peaks, only cytoarchitectonic areas available in the anatomy toolbox that had a >30% probability were considered. The cytoarchitectonic area with the highest probability was reported for each peak. When cytoarchitectonic areas with >30% probability were unavailable to determine a peak, the anatomical location of the peak was simply provided. In each cluster, we report peaks that were >8mm apart from each other in order of larger t-values. To facilitate visualization, we avoided reporting a peak for each cluster when it was identified in the cytoarchitectonic area or anatomical structure already reported for a peak with a higher t-value.

Abbreviation: PrG, precentral gyrus; SVC, small volume correction.
